# Pareto optimization of masked superstrings improves compression of pan-genome *k*-mer sets

**DOI:** 10.64898/2026.03.18.712440

**Authors:** Ján Plachý, Ondřej Sladký, Karel Břinda, Pavel Veselý

## Abstract

The growing interest in *k*-mer-based methods across bioinformatics calls for compact *k*-mer set representations that can be optimized for specific downstream applications. Recently, masked superstrings have provided such flexibility by moving beyond de Bruijn graph paths to general *k*-mer superstrings equipped with a binary mask, thereby subsuming Spectrum-Preserving String Sets and achieving compactness on arbitrary *k*-mer sets. However, existing methods optimize superstring length and mask properties in two separate steps, possibly missing solutions where a small increase in superstring length yields a substantial reduction in mask complexity.

Here, we introduce the first method for Pareto optimization of *k*-mer superstrings and masks, and apply it to the problem of compressing pan-genome *k*-mer sets. We model the compressibility of masked superstrings using an objective that combines superstring length and the number of runs in the mask. We prove that the resulting optimization problem is NP-hard and develop a heuristic based on iterative deepening search in the Aho-Corasick automaton. Using microbial pan-genome datasets, we characterize the Pareto front in the superstring-length/mask-run space and show that the front contains points that Pareto-dominate simplitigs and matchtigs. Finally, we demonstrate that Pareto-optimized masked superstrings improve pan-genome *k*-mer set compressibility by 12-19% when combined with neural-network compressors, achieving less than 1.2 bits per *k*-mer in common scenarios.

## INTRODUCTION

With the exponentially increasing amounts of sequenced data [1–3], *k*-mers and *k*-mer-based methods have become the workhorse of sequence bioinformatics. They have found their use across many applications including search in genomic data [3–6], metagenomic classification [7], and rapid diagnostics of antibiotic resistance [8,9], to name a few. Across all of them, the efficiency of *k*-mer data structures is tightly connected with the quality of the underlying *k*-mer set representations; for instance, it affects the time- and space-efficiency of *k*-mer indexes [10]. More compressed representations of *k*-mers allow these applications to process larger datasets on limited hardware, for instance, for serializing *k*-mers in large databases such as Logan [5]. This motivates the goal of designing representations that are highly compressible across a variety of use cases and can be optimized for specific downstream applications.

Recently, masked superstrings [11] have been proposed as a compressed representation for arbitrary *k*-mer sets, with the possibility to optimize them for specific applications. A masked superstring stores the *k*-mers as their superstring equipped with a binary mask, determining which of the superstring *k*-mers are represented. Masked superstrings generalize simplitigs/Spectrum-Preserving String Sets (SPSS) [12,13] and matchtigs [14], which are restricted to using overlaps of length *k* − 1 only. While computing the shortest superstring of *k*-mers is NP-hard, a greedy algorithm produces almost shortest superstrings in linear time [10,11]. Greedy masked superstrings can be slightly better compressed than SPSS-based methods [11] and representations based on them have found applications in space-efficient *k*-mer set indexing [10] and in data structures for performing set operations with *k*-mers [15].

However, all construction methods for masked superstrings, simplitigs, or matchtigs focus on the total representation length as the primary size measure, which fails to capture nuanced compressibility tradeoffs between superstrings of different lengths and their masks. For instance, while simplitigs and matchtigs provide suboptimal superstrings, their masks, corresponding to sequence delimiters, tend to be highly compressible as they typically contain only a few runs of consecutive ones. On the other hand, greedy masked superstrings require masks with potentially more complex and less compressible structure, resulting in comparative overheads that are not captured by the representation length.

Here, we provide the first algorithm for constructing a textual representation of *k*-mer sets that optimizes for the structure of the representation beyond length. Building upon the framework of masked superstrings [11], we focus on jointly optimizing a linear function of the superstring length and the number of runs in the mask as a proxy for mask compressibility. We show that for any linear combination, the objective is NP-hard and develop a heuristic algorithm for Pareto optimization based on iterative deepening search in the Aho-Corasick automaton. Using microbial pan-genome *k*-mer sets as examples, we study the Pareto front and show that with appropriate scaling between superstring length and the number of runs, microbial pan-genomic *k*-mer sets are close to lower bounds on the superstring length or on the number of runs. Finally, we demonstrate that the Pareto-optimized masked superstrings improve disk compression of *k*-mer sets with classical dictionary-based as well as specialized neural network-based compressors.

## PRELIMINARIES

### Spectrum-Preserving String Sets (SPSS)

Simplitigs/spectrum-preserving string sets [12,13] are vertex-disjoint paths in the de Bruijn graph of a given *k*-mer set *K*, a generalization of unitigs without the restriction of stopping at branching vertices. Eulertigs [16] are the length-optimal variant of simplitigs. Matchtigs [14] additionally allow any *k*-mer of *K* to appear multiple times in the strings, and can be computed in polynomial time. For simplicity, we refer to all these representations as SPSS.

### Masked superstrings (MS) of *k*-mers

The masked superstring (MS) [11] of a *k*-mer set *K* is a representation which allows “false positive” *k*-mers, i.e., those not present in *K* may appear in the superstring. Formally, MS is a pair (*S*, *M*) such that: (i) *S* is a superstring containing all *k*-mers of *K* as substrings, and (ii) *M* is a binary mask that masks out the false positive *k*-mers. The mask is of the same length as the superstring and has the property that each *k*-mer from *K* has at least one occurrence *S* [*i*:*i* + *k* − 1] such that *M* [*i*] = 1, whereas the “false positive” *k*-mers outside *K* have all such *M* [*i*] = 0. As only one occurrence is required to be masked with 1, this means that potentially multiple masks represent the same *k*-mer set *K* with a fixed superstring *S*. This leaves room for further mask optimizations. Masked superstrings generalize SPSS as for any SPSS representation of *K*, MS can be constructed by concatenating all sequences *S*_1_… *S*_*n*_(simplitigs or matchtigs) into a single superstring *S* and concatenating a sequence of ones of length | *S*_*i*_| − (*k* − 1), interspersed by a sequence of *k* − 1 zeros. Exactly those MS where all runs of zeros in the mask are of length *k* − 1 have an equivalent SPSS representation of the same length. Similarly to SPSS, which are formed by paths in the de Bruijn graph, a superstring corresponds to a Hamiltonian path in the *overlap graph*, where vertices are *k*-mers and there is a directed edge between each pair of *k*-mers, weighted by their overlap.

### State-of-the-art masked superstring construction algorithms

The only existing methods for MS construction start by aiming for the smallest possible length, either through SPSS algorithms [12 –14,16] or directly greedily approximating the shortest superstring [11]. In [11] this first step is followed by a second step, where superstring *S* is fixed and mask *M* is optimized with respect to a specified criterion; for instance, to minimize or maximize the number of ones or to minimize the number of runs of ones, denoted by *runs*_1_(*M*), where a run is a maximal substring of *M* of the given symbol. Minimization and maximization of ones can be performed simply in linear time, while minimization of the number of runs is NP-hard [11]. While this two-step process can be efficiently implemented, its disadvantage is that the initial computation of the superstring, focusing solely on its length, may restrict the space of admissible masks so that none has good compressibility properties.

### Aho-Corasick automaton

The Aho-Corasick automaton (AC) for a *k*-mer set *K* consists of the prefix tree for the *k*-mers and failure links, leading from a prefix to its longest proper suffix that is a node in the tree. The edges of the prefix tree are called forward links. The depth of every node is the length of the prefix it represents. We will call a set of all nodes of certain depth *d* the *d*-th level of AC. In the context of this work, AC has been applied for implementing greedy superstring computation in linear time [17] or for representing overlap graphs in linear space, i.e., constructing so-called *hierarchical overlap graphs* (HOG) [18,19].

## METHODS

### Pareto optimization of superstrings and masks

Our aim is to compute *k*-mer set textual representations with improved compressibility properties. However, compressibility is a very complex measure, with many possible ways to express it. We propose modeling it as a combination of superstring length and the number of runs in the mask, weighted by a *run penalty* parameter *P*. That is, we study the Pareto optimization problem of computing a masked superstring (*S*, *M*) minimizing

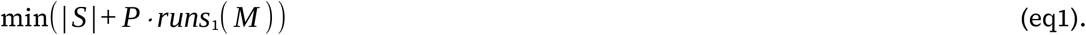

This directly corresponds to storing the superstring in a plain 2-bit encoding and the mask in a run-length encoding, similarly to how these representations are stored within modern *k*-mer indexes [10,20,21]. More generally, the superstring length is a proxy to its compressibility as the superstring is typically less compressible than the mask. Similarly, the number of runs in the mask is a proxy for mask compression as with very few runs, run-length encoding yields a much smaller description. However, note that the superstring length and the number of runs can be in opposition as minimizing one might increase the other (**Fig. 1b**).

**Fig 1.**
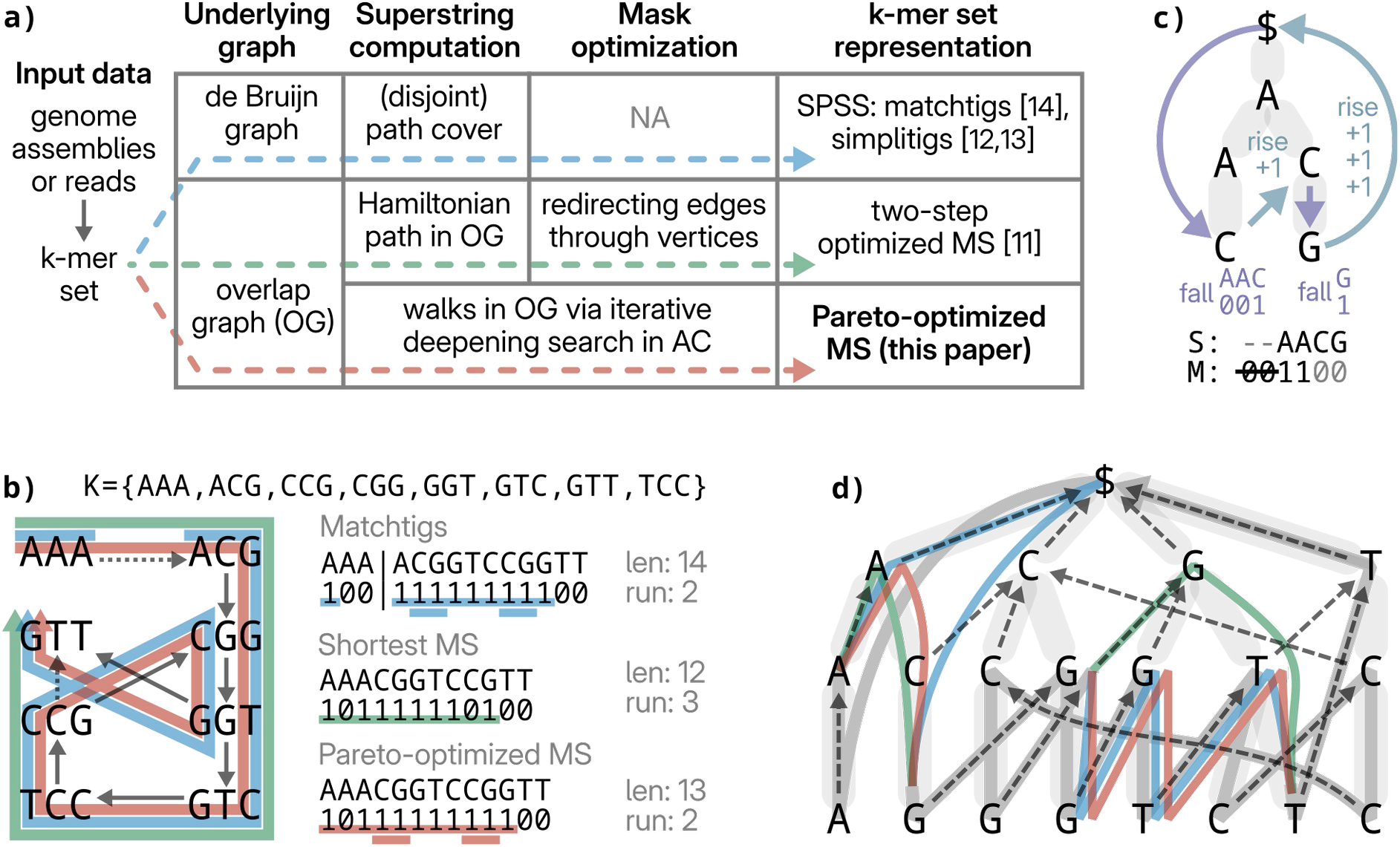
Overview of our approach. **a)** The computation pipeline for SPSS, MS with two step optimization, and MS with Pareto optimization. All methods compute different coverings of *k*-mer graphs. While SPSS and the two-step method first aim to compute the shortest representation and mask optimization is either missing or secondary, Pareto optimization computes walks that simultaneously optimize both the superstring and the mask. **b)** An example of three representations for *k*-mer set *K*, namely shortest matchtigs, shortest MS, and MS constructed by Pareto optimization (with *P≥* 2). Overlaps of length *k* − 1 are represented by solid arrows and selected overlaps of length *k* − 2 by dotted arrows. Observe that unlike MS-based methods, matchtigs cannot utilize the overlap of length 1 < *k* − 1 from AAA to ACG, increasing their total length, and that Pareto optimization reuses CGG and GGT, saving one run for a slight increase in total length. **c)** An illustration of *fall* and *rise*, two elementary operations on a small AC automaton along with the symbols emitted by fall and penalty incurred by rise. For simplicity, each level penalty is 1. The emitted superstring is a concatenation of the emitted symbols. The emitted mask is a concatenation of emitted mask symbols, ignoring the initial *k* − 1 zeros and adding trailing *k* − 1 zeros. **d)** The AC automaton of *k*-mer set *K*, with walks corresponding to the representations from b); the dark grey walk is common to all three representations. Light gray bars represent forward links, dashed arrows represent failure links. Matchtigs are restricted to the two bottommost levels and the root, with additional ascent to the root not done by MS-based methods. The path CG → CGG → GG → GGT → GT is used twice by both matchtigs and Pareto optimization while the MS of the optimal length ascends to G at depth *k* − 2, starting a new run in the mask.

The Pareto optimization problem is in general NP-hard, as even the shortest superstring problem for *k*-mers is NP-hard [11]. Moreover, as we show in Theorem 1, for any run penalty *P*, finding the optimal value of the objective remains NP-hard; the proof is in **Note S1**.

#### Theorem 1 (NP-hardness of Pareto optimization)

*For any constant P* > 0 *and any value of k* > 4 *⋅* log_4_(| *K* |)+ 5, *where K is a set of k-mers, computing an optimal MS of K according to the objective function* | *S* |+ *P runs*_1_(*M*) *is NP-hard*.

### Aho-Corasick-based reformulation of SPSS and MS constructions

Since the problem of Pareto optimization is NP-hard, we resort to searching for near-optimal heuristics. However, before we describe the heuristic, we first need to set up a different view on algorithms for masked superstring construction. We design two low-level operations using the Aho-Corasick tree of *k*-mers, *fall* and *rise*, and show that existing algorithms can be viewed in this new framework as performing these two low-level operations. We design our heuristic for Pareto optimization in the next section.

Consider the Aho-Corasick (AC) automaton of the *k*-mer set. For each level *L* from 1 to *k*, we define *level penalty π*_L_based on the optimization problem. Our aim is to construct a walk in the AC automaton and emit characters of a masked superstring during the walk. We use two elementary operations, defined for a node *v* of the automaton as follows:

### Fall

Descend to a leaf in the subtree of *v* via forward edges. All characters on forward edges are *emitted* in the top-down order. In addition, for each such edge, a mask symbol is emitted: 0 for inner nodes, and 1 for the final step to the leaf. Fall does not pay any penalty.

### Rise

Ascend to a node on a higher level via the failure link. A cumulative penalty for each traversed level is paid. More formally, if the failure link goes from level *i* to level *j* < *i*, then the penalty *π*_ij_= *π*_i_+ *π*_i-1_+ … + *π*_j+1_is paid. Rise does not emit any characters.

In this framework, the goal is to construct, using these two operations, a *closed covering walk* that starts and ends in the root of AC and covers all leaves, which correspond to the individual *k*-mers. We say that a closed covering walk emits a superstring *S*, which is simply the concatenation of characters emitted by the walk. However, the closed walk emits a sequence of mask symbols corresponding to a slightly different formulation of the mask, where a *k*-mer is masked at its *last* character, instead of the first. This leads to a redundant prefix of *k* − 1 zeros and a missing suffix of *k* − 1 zeros. We thus apply a postprocessing step on the mask: From the concatenation of mask symbols emitted, we ignore the *k* − 1 leading zeros and append *k* − 1 trailing zeros. See **Fig. 1c** for an example of rise and fall operations.

#### Theorem 2 (Closed covering walks correspond to masked superstrings)

*Given an AC automaton of a k-mer set K, each closed covering walk emits a masked superstring representing K. Each masked superstring with M* [0] = 1 *and no runs of zeros longer than k* − 1 *has a unique closed covering walk spelling it*.

**Proof of Theorem 2**: To see that a closed covering walk emits an MS of *K*, observe that whenever we are at a leaf, the last *k* characters of the so far emitted superstring are the *k*-mer corresponding to the leaf and the last mask symbol is 1, implying that after the postprocessing of the emitted mask, the *k*-mer is correctly masked as represented by the masked superstring. Meanwhile, a *k*-mer outside of *K* cannot be masked as represented; indeed, its occurrences must be masked with 0s as we emit 1 only in leaves, which correspond to *k*-mers from *K*. We prove the other part by a construction of the walk. We start in the root and we read the given MS. For each 1 in *M*, we descend to the leaf for the current *k*-mer of the MS, and we also ascend by one level, moving by one character in the superstring as well as the mask. If the 1 in *M* is followed by a run of zeros, we ascend by that many levels and move by the same number of characters in the masked superstring. Before the next 1 in *M*, we are guaranteed to be at a prefix of the current *k*-mer, so the ascend made corresponds to the rise operations and the fall operation is possible from the current node. In addition, each *k*-mer of *K* is visited as it is masked by the MS, thus the walk is covering. The last run of *k* − 1 zeros guarantees that we end up in the root, implying the walk is closed. The uniqueness part follows from the fact that two distinct walks emit different masked superstrings. **Q.E.D**.

The restriction to masked superstrings with masks starting with 1 and not containing a run of zeros longer than *k* − 1 is necessary, since the superstring positions that are not part of any represented *k*-mer may contain arbitrary sequences outside of the represented *k*-mer set. This is however not an issue from the practical side as these positions can be removed without modifying the underlying set and without worsening any practically relevant measures. For an illustration of correspondence of walks in an AC automaton to representations, see **Fig. 1d**.

Naturally, the problem is to find a closed covering walk in the AC automaton that minimizes the total penalty of its rise operations. Depending on the level penalties chosen, the optimal walk corresponds to different representations of *k*-mer sets. The different level penalties needed for known representations are given in the following theorems; see also **Table S1**.

#### Theorem 3 (Shortest superstrings)

*Given a k-mer set K and its AC automaton, and considering level penalty π*_*L*_ = 1 *for all levels L, the optimal closed covering walk of the automaton spells the shortest superstring of K*.

**Proof:** For every letter emitted, one level is descended, and for every level ascended, penalty of 1 is paid. Hence, the total penalty is the length of the superstring and since closed covering walks and masked superstrings are in one-to-one correspondence by Theorem 2, the optimal closed covering walk corresponds to the shortest superstring. **Q.E.D**.

#### Theorem 4 (Matchtigs)

*Given a k-mer set K and its AC automaton, and considering level penalty π*_*k*_ = 1, *π*_*k-1*_ = 2 *k* − 3, *and π*_*L*_ = − 1 *for all levels L* < *k* − 1, *the optimal closed covering walk of the automaton emits matchtigs of K minimizing the total length*.

**Proof**: First, observe that whenever the optimal walk rises above the level *k* − 1, it rises all the way up to the root, due to the penalties of the upper levels being negative. Right afterward the walk gets to the root, it has to fall to a leaf, emitting exactly *k* − 1 zeros in the mask. There is no other possibility to emit zeros, thus each run of zeros has length exactly *k* − 1, implying that the emitted sequence corresponds to matchtigs. For the sequence of rise steps to the root, the cumulative penalty is 1 + 2 *k* − 3 − (*k* − 2)= *k*. On the other hand, if the walk rises to level *k* − 1 and then immediately falls to a leaf, it pays a penalty of 1. Consider a particular matchtig emitted: rising to the root happens exactly once and rising to level *k* − 1 happens once for every *k*-mer except for the last one. Hence, the penalty corresponding to the matchtig is equal to the length of the matchtig. Therefore, the total penalty is the length of the emitted matchtigs and the optimal walk emits shortest matchtigs. **Q.E.D**.

#### Theorem 5 (Simplitigs)

*Given a k-mer set K and its AC automaton, and considering level penalty π*_*k*_ = 1, *π*_*k-1*_ = *k* − 3 / 2, *and π*_*L*_ = − 1 *for all levels L* < *k* − 1, *the optimal closed covering walk of the automaton emits simplitigs of K of shortest possible length*.

**Proof**: Using the same reasoning as in Theorem 4, the negative penalties guarantee that each run of zeros in the optimal walk has length *k* − 1, meaning the optimal walk corresponds to an SPSS. Next, we prove by contradiction that in the optimal walk, no *k*-mer appears twice with its mask symbol set to 1. Suppose this were the case, and there is a *k*-mer *Q* with more than one masked occurrence. If at least one of the occurrences of *Q* was at the beginning or at the end of a run of ones, the leaf corresponding to the occurrence of *Q* can be removed from the walk, decreasing the penalty by 1 without violating the covering property, contradicting optimality. If it is in the middle of a run, the run can be split on this *k*-mer, by rising to the root instead of falling to the leaf *Q* and then falling to the subsequent leaf; this saves a penalty of 1 and incurs cost *k* − 3 / 2 − (*k* − 2)= 1 / 2, hence saving 1 / 2. Therefore each leaf appears exactly once in the walk, implying that the superstring corresponds to simplitigs. Moreover, the total penalty is | *K* |+ # *simplitigs* / 2. As minimizing the number of simplitigs is equivalent to minimizing their total length, the walk emits optimal simplitigs. **Q.E.D**.

In addition to existing representations, we can easily formulate the Pareto optimization problem in this reformulation by setting appropriate level penalties.

#### Theorem 6 (Pareto optimization)

*Given a k-mer set K and its AC automaton and considering a level penalty π*_*k-1*_ = *P* + 1 *and π*_*L*_ = 1 *for all levels L≠ k* − 1, *the optimal closed covering walk of the automaton emits a masked superstring* (*S*, *M*) *minimizing (eq1), that is* | *S* |+ *P runs*_*1*_(*M*).

**Proof**: Similarly as in Theorem 3, each level has penalty of at least one, thus we pay one for each character of the superstring. Only level *k* − 1 has an additional penalty of *P*. Since between every two runs of ones and also after the last run of ones, level *k* − 1 is crossed once, this extra penalty contributes *P · runs*_*1*_(*M*) in total. Hence, the total penalty is exactly | *S* |+ *P · uns*_*1*_(*M*) and the optimal closed covering walk emits the optimal MS with respect to (eq1). **Q.E.D**.

### Greedy for Pareto optimization of superstring length and the number of runs

We utilize the reformulation of optimal masked superstrings for (eq1) given by Theorems 2 and 6 as an optimal closed covering walk of the AC automaton with the level penalties *π*_k-1_= *P* + 1, and *π*_L_= 1 for all levels *L ≠ k* − 1. However, the problem is NP-hard by Theorem 1, and so we apply a greedy heuristic.

At each step, we identify the two not-yet connected leaves where the walk from one to the other incurs the smallest penalty and connect them. This may not produce optimal MS w.r.t. (eq1) as it may be better to perform a locally worse step that improves the penalty later on.

To determine the pair with the shortest subpath between them, we perform iterative deepening depth-first search (DFS) on the nodes of the automaton. In more detail, we perform iterations with a maximum subwalk penalty starting from 1 in the first iteration and finishing at *P* + *k*. In each iteration, for each leaf with no outgoing subwalk, we sequentially perform DFS, terminating the search once the penalty exceeds the maximum subwalk penalty for this iteration. Once a different leaf with no incoming subwalk is reached, the two leaves are connected via a subwalk. Due to the nature of iterative deepening, whenever two *k*-mers are connected, there is no pair with strictly smaller penalty for their subwalk. For the very last connection, we ensure that this connection goes through the root – this will be the start and end of the walk. As the upper bound for the penalty between two leaves is *P* + *k* for walking up to the root and down, after *P* + *k* iterations, all leaves are connected.

### Algorithmic optimizations

The main limitation of using an AC automaton is the need to store *O* (| *K* | *⋅ k*) nodes and edges, and we thus do not store it explicitly. Instead, we only store the *k*-mers of *K* in the lexicographic order. Each AC node is represented as a pair of index and depth. The index points to the first *k*-mer in the lexicographically sorted set *K* with the given node as a prefix, and the depth is the length of the prefix. To support the fall operation, we find all leaves with the prefix of a given node, by starting at the node’s index and traversing the sorted *K* while the prefixes match. For the rise operation, to find whether a suffix of a given length of a node exists, we binary search the sorted *K*. To speed up the search, the first index is stored for every possible log_4_(| *K* |)-character prefix, requiring additional constant space per k-mer.

Next, we reduce the number of repetitive operations performed by the algorithm. We define a *subsearch* of a given search in the AC automaton as a pair of its starting node and an amount of *remaining penalty* not yet spent in the search before the start of the subsearch. We notice that repeating any subsearch finds no leaf to connect if the former search found no leaf, as in later phases of the algorithm, the number of suitable leaves is low. Thus for every leaf, we store the maximum remaining penalty it was visited with, and we only extend the search if we are able to assign a strictly higher remaining penalty. This slightly decreases the quality of the solutions, but improves the practical running time, as no leaf is used to extend the search more than *P* + *k* times in total, and the total number of rise and fall operations is *O* (| *K* | *k* (*k* + *P*)). However, every fall operation may yield up to *O* (| *K* |) leaves to explore, resulting in a worst case time complexity *O* (| *K* |^2^*k* (*k* + *P*)). Nevertheless, on a typical input, the search finishes successfully in the first iteration for most of the leaves, and thus fall visits only a few leaves, meaning that the typical real time complexity is between the worst case and best case complexity of *O* (| *K* | *k*).

To prevent creating cycles between the leaves being connected, we use a union-find data structure with path compression and asymmetric unions. Finally, reverse complements are handled by adding them to the set as a first step, and performing all union operations on both the paths and their reverse complements as well. The pseudocode of the implementation is given in **Alg. S1**.

### Computing the lower bound on the number of runs

To evaluate the Pareto optimization algorithm, we seek to get lower bounds on the objective function (eq1). A non-tight lower bound for the superstring length is given by the shortest cycle cover of the corresponding overlap graph [22], which can be computed in linear time. Next, consider the minimum number of runs in a mask for any superstring. While this problem is NP-hard for a fixed superstring [11], we show that if the superstring is not fixed, the minimum number of runs can be computed in polynomial time by reducing the problem to finding the minimum number of matchtigs representing the same set; as the next lemma shows, these two numbers are equal (the proof is in **Note S2**).

#### Lemma 1.

*Given a k-mer set K, the minimum number of runs in the mask of any MS representing K is exactly equal to the minimum number of matchtigs needed to represent K*.

Note that the minimum possible number of matchtigs is not equal to the number of typically computed matchtigs, as tools for matchtigs computation optimize for length and not for count.

The lower bound on the number of runs is computed as follows: In the de Bruijn graph of *K*, each strongly connected component can be covered by a single matchtig that can enter and leave the component at any *k*-mer. Hence, we contract the strongly connected components, using the standard linear time algorithm. This reduces the task to finding the minimum number of paths to cover the directed acyclic graph (DAG) of strongly connected components [23], which is reduced to an instance of the maximum bipartite matching [24]. This can be solved using a standard algorithm such as the Kuhn’s algorithm, which is practically efficient. In **Note S2**, we additionally discuss how to deal with reverse complements.

### Experimental evaluation

We implemented the proposed Pareto optimization algorithm in C++ using the internals and interface of KmerCamel [11], and made it available in the supplementary repository^1^.

We evaluated the Pareto optimization algorithm using *k*-mer sets obtained from pan-genome datasets of various sizes, including *S. pneumoniae* with 616 genomes and 9 million 31-mers, SARS-CoV-2 with 16 million genomes and 12 million 31-mers, and *E. coli* with 85,885 genomes and 450 million 31-mers, from [25]. We compared it to the previous state-of-the-art representations, namely greedy matchtigs [14] (computing matchtigs of optimal length took too much time for larger datasets or for *k* = 15), greedy masked superstrings with mask optimized using one of three objectives (min-one and max-one, minimizing and maximizing the number of ones in the mask, respectively, and min-run, minimizing the number of runs of ones in the mask) [11], and eulertigs (simplitigs of minimal length) [16]. For comparison, we also included commonly used unitigs (**Supplementary repository^1^**); nevertheless, we found that on our datasets they consistently performed poorly in both target criteria. Additionally, we computed lower bounds for the superstring length [11] and the smallest number of runs of ones in the mask of any masked superstring representation.

The experiments were implemented using a Snakemake pipeline, which first computed the masked superstring representations, then compressed them using several presets (using xz, bzip2, and GeCo3 [26], a neural network-based DNA compressor), and, finally, analyzed the properties of superstrings, masks, and the compressed files. Furthermore, we verified correctness of our pipeline, i.e., that the decompressed *k*-mer set remains the same as the original *k*-mer set (**Note S3**). As a baseline for overall effectiveness of our approach, we used the ESSCompress tool [27] (v3.1), which computes its own compressed representation. The pipeline alongside the results is provided in the **Supplementary repository^1^**.

The indicative resources needed for computation for SARS-CoV-2 pan-genome dataset and specifications of hardware used for the benchmarking are provided in Table S2.

### Pareto optimization dominates known methods

First, we asked about the effect of varying the *run penalty* (*P*) parameter on the properties of the resulting masked superstrings. We found (Fig. 2) that heuristic Pareto optimization provides a substantially better characterization of the Pareto front between the superstring length and the number of runs of ones in the mask, compared to state-of-the-art *k*-mer set representation methods. As expected, with increasing values of *P*, the length increases and the number of runs decreases, since the greedy heuristic may decide to construct a longer superstring to save more runs of ones.

**Fig 2.**
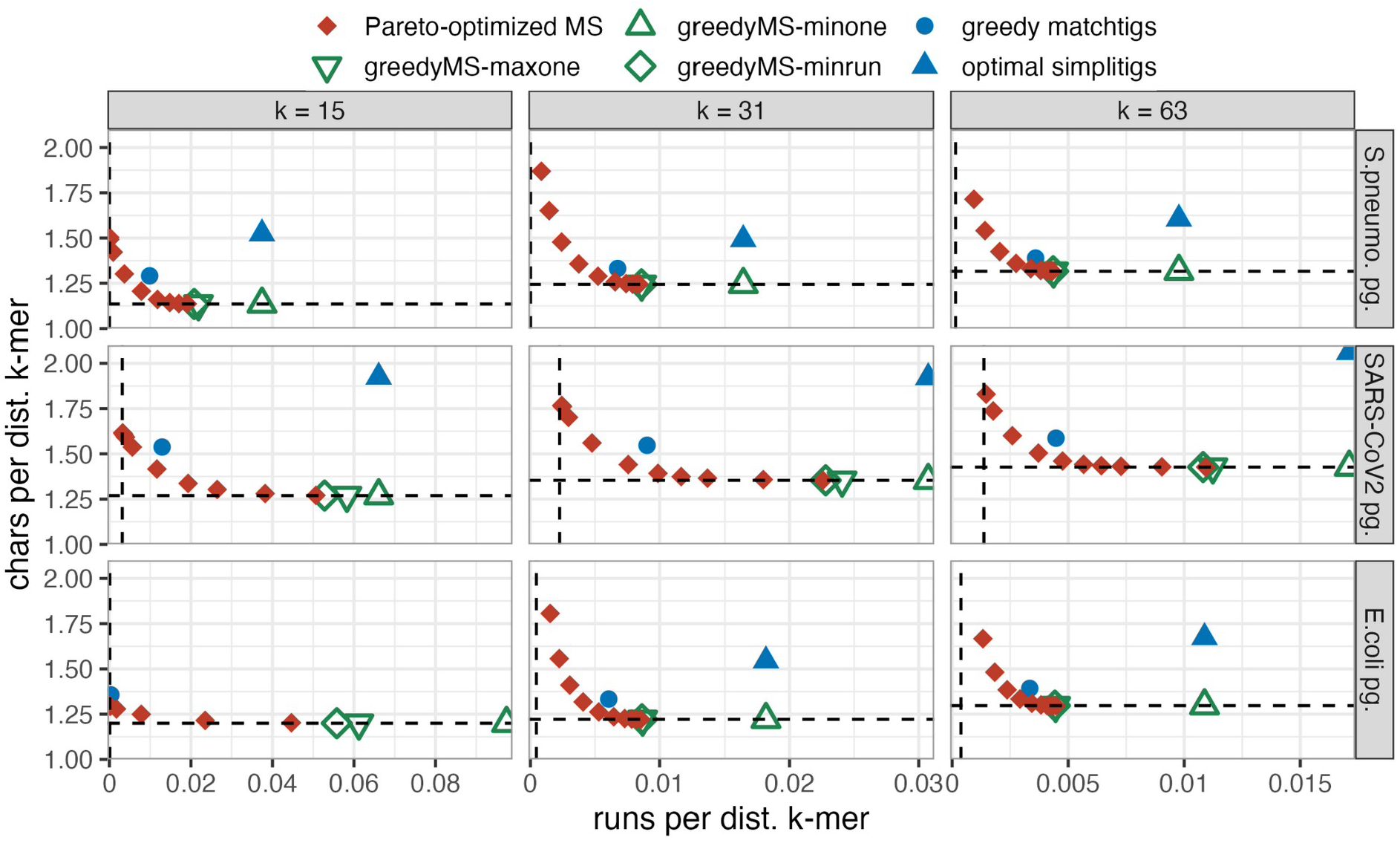
Tradeoffs between the superstring length and number of runs of ones, including the Pareto front. For selected microbial pan-genomes and values of *k*, we display the superstring length per (distinct canonical) *k*-mer on the *y*-axis, and the number of runs of ones in the mask per *k*-mer on the *x*-axis. The dashed horizontal line depicts the lower bound on the superstring length, and the dashed vertical line is the lower bound on the number of runs of ones in any possible mask. The Pareto-optimal superstrings are guaranteed to lie between Pareto optimization (red points) and dashed lower bounds.

We first compare the results of Pareto optimization to greedy masked superstrings, which are within 0.01 % of the lower bound on the superstring length in all experiments. First, we consider *k* = 31 and *k* = 63. For small values of *P≤* 8, the length increase of the superstring for most of the datasets stays within 1 % of the lower bound; however, the number of runs drops significantly, by up to 10 % for *k* = 31 and 5 % for *k* = 63. For the SARS-CoV-2 pan-genome, which consists of a large number of short genomes, the length increase is *≤* 3 %, while the number of runs drops by up to 50 % and 30 %, respectively.

For larger values of *P*, such as *P* = 32, the length of the superstring increases by *≤* 6 %, while the number of runs drops by *≥* 20 %, or 2-3 times for the SARS-CoV-2 dataset. For even larger values of *P* (up to 512), the length further increases by up to 50-75%, while the number of runs slowly approaches the lower bound; in some cases, the lower bound is almost reached and increasing *P* over a certain threshold yields almost no changes, while for instance for the *E. coli* pangenome, an even higher value of *P* would probably be needed to reach the lower bound.

When considering short *k*-mers with *k* = 15, the optimization behaves similarly as for the larger ones, except that Pareto optimized MS get to the lower bound faster with increasing *P*. For *S. pneumoniae* pan-genome, even with a small *P* = 8, the length increases by *≤* 6 %, while the number of runs drops by 45-50%. For the SARS-CoV-2 pan-genome and *P* = 8, the number of runs drops by 64 %, decreasing almost three times compared to the mask optimized with the min-run criterion but using a fixed superstring. For the *E. coli* pan-genome, which contains nearly 190 million 15-mers, the number of runs drops about 25 times for *P* = 8 and about 380 times for *P* = 32, getting within 25 % of the lower bound.

Greedy matchtigs are Pareto-dominated by Pareto-optimized MS. When considering either the superstring length or the number of runs in the mask, we are able to find a value of *P* that is not worse in one criterion, while in the other criterion, it outperforms the matchtigs by 30 - 50 % of a difference to the respective lower bound. Simplitigs, even optimal, are far longer and also have more runs than matchtigs or greedy MS with max-one or min-run mask optimization.

### Pareto optimization improves compressibility of textual *k*-mer set representations

Next, we sought to understand the relationship between the *run penalty* parameter *P* and the actual compressibility of the masked superstrings. We asked about two distinct practical use cases: disk compression (for long-term storage) and in-memory compression (allowing rapid access to the data). We measured the results for Pareto optimization with several chosen values of *P* between 1 and 512, and also for simplitigs, greedy matchtigs, and the masked superstrings with min-run and max-one mask optimization criteria.

For disk compression, we measured the compressibility of the superstring and the mask separately, using the size of the resulting files after compression with several protocols, as explained in **Note S3**. Surprisingly, we found (**Fig. 3**) that even if the superstring length increases significantly for large values of *P*, such as 256 or 512 (as reported in **Fig. 2**), the compression is more effective for the superstring itself than for smaller values of *P* when using protocol 4 (the specialized GeCo3 tool with parameter setup for maximal compression). As expected, the compressibility of masks improves with increasing run penalty *P* and decreasing number of runs of ones in the mask. The Pareto optimization outperforms both greedy MS and matchtigs by 12 - 19 % for *k* = 31. Using an optimized compression protocol with GeCo3 (protocol 4 in **Note S3**), we were able to represent the 31-mer sets of the *S. pneumoniae* and *E. coli* pan-genomes in slightly less than 1.2 bits per *k*-mer and SARS-CoV-2 in less than 1.1 bits per *k*-mer. For *k* = 63, we obtained approximately 0.8 bits per *k*-mer for bacterial pan-genomes and even 0.6 bits per *k*-mer for SARS-CoV-2. Results for standard compression and for *k* = 63 are provided in **Fig. S1** and **Fig. S2**.

**Fig 3.**
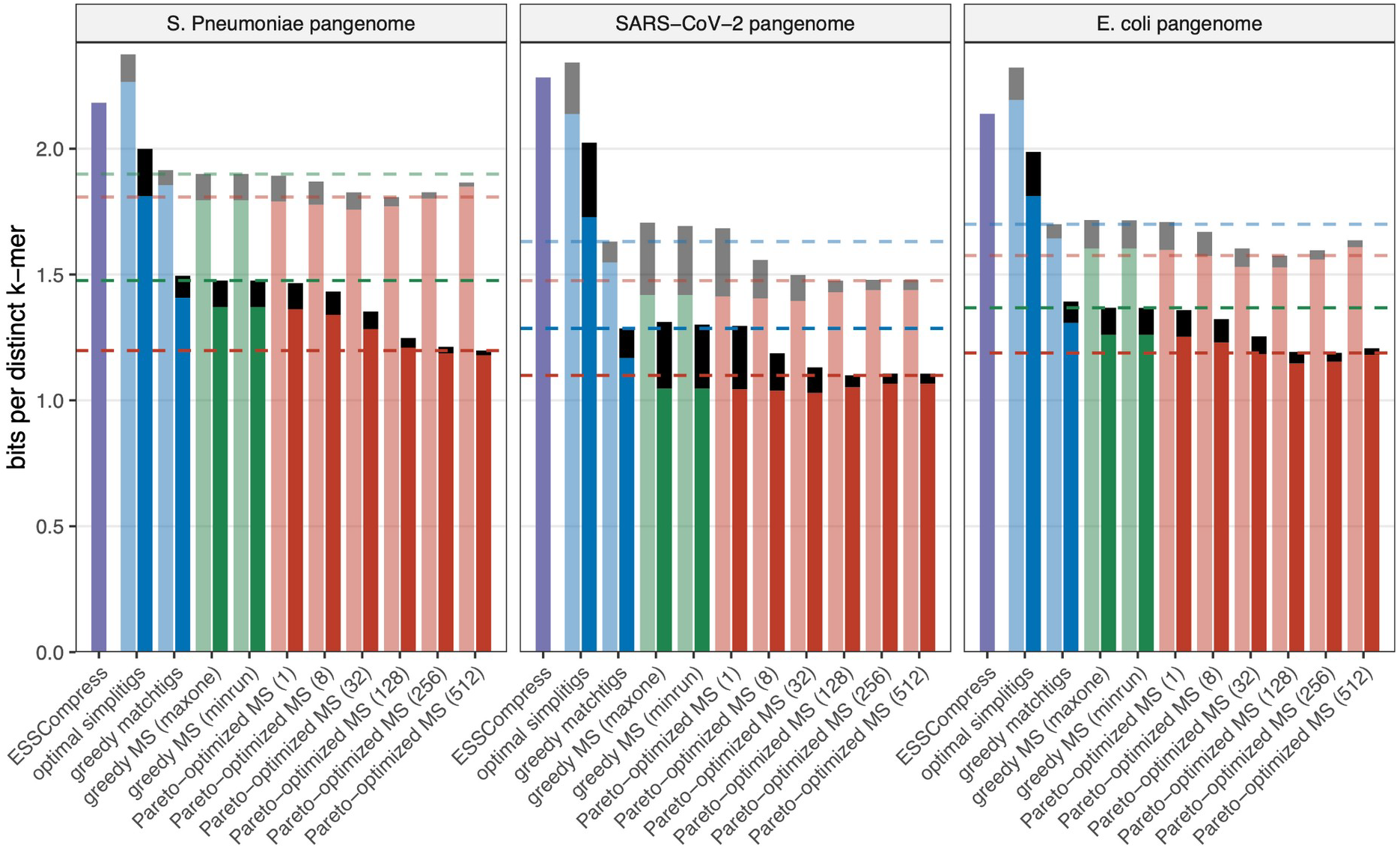
Disk compression of masked superstring representations for selected datasets and *k* = 31 using xz and GeCo3. For selected methods and parameters, we display the compressed sizes of superstrings (coloured bars starting at zero) and masks (black bars on top of the superstring bars). The bar for ESSCompress represents the total size of the *k*-mer set compressed with ESSCompress. Opaque bars represent compression with GeCo3, while semi-transparent bars represent compression with xz for superstrings and bzip2 for masks; all tools with maximal compression presets of parameters **(Note S3)**. The green or blue dashed line represents the best compressed size of the state-of-the-art representation (masked superstrings with min-run mask optimization or matchtigs); the red dashed line represents the best compressed size achieved using Pareto optimization.

For in-memory compression, the superstring was encoded using two bits per character (no compression), while the mask was compressed using Elias-Fano encoding [28,29]. Results for *k* = 15 and *k* = 31 are provided in **Fig. S3**. For *k* = 31, the improvements in the in-memory compression are negligible, the best one being 3.4 % for SARS-CoV-2 pan-genome. For *k* = 15, the improvements are between 2 % and 5 %. The main reason for such small improvement is that the length of the superstring, which is nearly optimal for greedy MS, accounts for most of the in-memory size of the MS. However, the compression of the mask is improved by more than 60% when compared to matchtigs and by more than 70% when compared to greedily constructed masked superstrings. In conclusion, even though the in-memory compressibility of the mask is greatly improved, storing the superstring in the plain 2-bit encoding still vastly dominates space requirements.

## DISCUSSION

In this work, we designed the first algorithm for optimizing textual *k*-mer set representations that goes beyond minimizing length. Our algorithm Pareto-optimizes both the superstring length and the number of runs in the mask. We confirmed our initial hypothesis that the resulting masks are substantially more compressible, and surprisingly found that the masked superstring itself becomes – despite being longer – more compressible than length-optimized MS and matchtigs. Our hypothesis for this intriguing phenomenon is that a high penalty for a new run diminishes the number of distinct *k*-mers in the superstring and leads to a better preservation of the original statistical biases (e.g., biases related to replication origin). Overall, our pipeline achieves lossless compression using less than 1.2 bits per *k*-mer for the considered pan-genomes and *k* = 31, while for *k* = 63, we compressed the *k*-mer sets to even less than 0.9 bits per *k*-mer.

Our Pareto optimization heuristic trades increased construction time for improved compressibility. The main current limitation, compared to the state-of-the-art tools for SPSS or MS computation, is longer construction time, namely, on SARS-CoV-2 pan-genome our algorithm is 5 - 10 times slower (and more for higher run penalty). Nevertheless, additional acceleration and scalability may be achieved via low-level optimization, parallelization, and possibly also in combination with succinct AC representation techniques [30].

Overall, our results showcase the strong compressive properties of masked superstrings as a representation of *k*-mer sets. Furthermore, our work opens a new direction for application-specific optimization of *k*-mer set representations.

## Acknowledgements

This work was supported by the French National Research Agency (ANR-24-CE45-1226; REALL); Czech Science Foundation (24-10306S); Czech Ministry of Education, Youth and Sports (ERC-CZ LL2406); Charles University (UNCE 24/SCI/008); French Ministry for Europe and Foreign Affairs, French Ministry for Higher Education and Research, and Czech Ministry for Education, Youth and Sports (PHC BARRANDE 2025 grant no. 52374TC; EFFIMAS). Computational resources were provided by the e-INFRA CZ project (ID:90254), supported by the Ministry of Education, Youth and Sports of the Czech Republic.

1 https://github.com/Jajopi/ms-pareto-optimization-supplement

