## Supplement for "Pareto optimization of masked superstrings improves compression of pan-genome *k*-mer sets"

### Supplementary figures

#### Figure S1: Maximal disk compression for k = 63

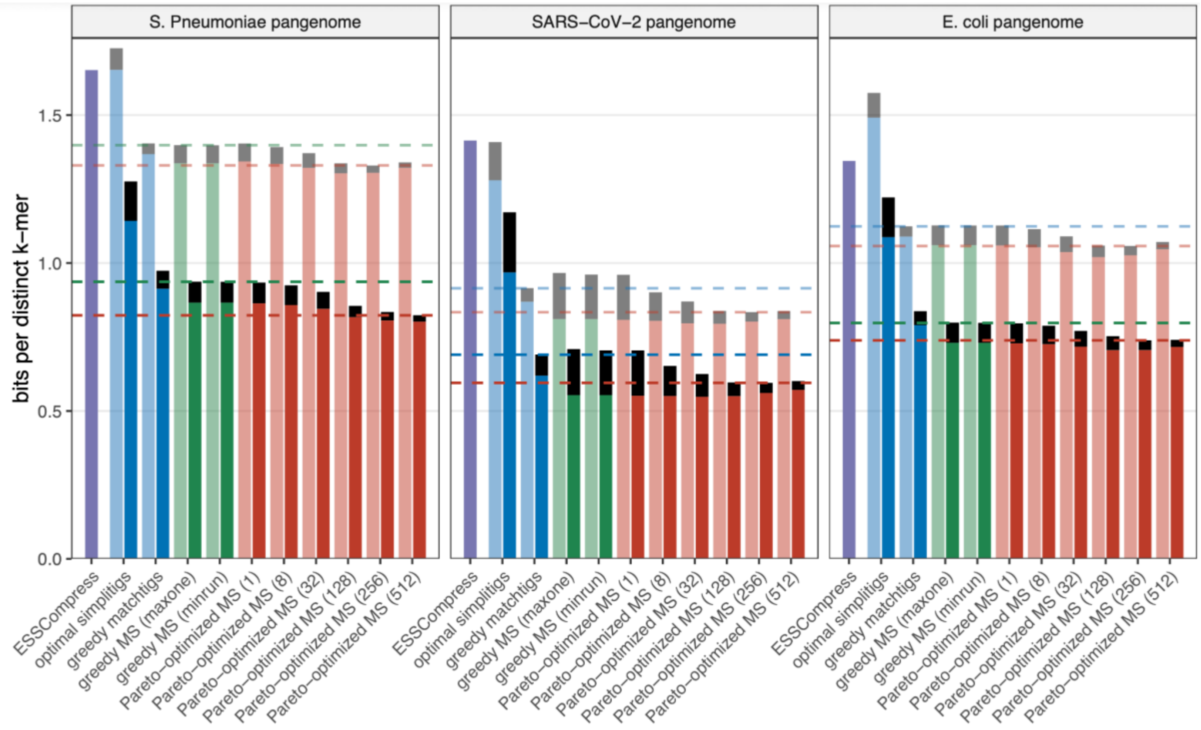

**Fig. S1. Disk compression of MS representations for selected datasets and *k = 63*, using maximal compression setup.**

For selected methods and parameters, we display the compressed sizes of superstrings (coloured bars starting at zero) and masks (black bars starting at the top of superstring ones). The bar for ESSCompress represents the total size of the *k*-mer set compressed with ESSCompress. Opaque bars represent compression with GeCo3, while semi-transparent bars represent compression with xz for superstrings and bzip2 for masks. The parameters for these tools are specified in presets 2 and 4 in **Note S3**. The green or blue dashed line represents the best compressed size of the state-of-the-art representation (masked superstrings with min-run mask optimization or matchtigs), and the red dashed line represents the best compressed size achieved using Pareto optimization.

### Figure S2: Standard disk compression, $k = 31$ and $k = 63$

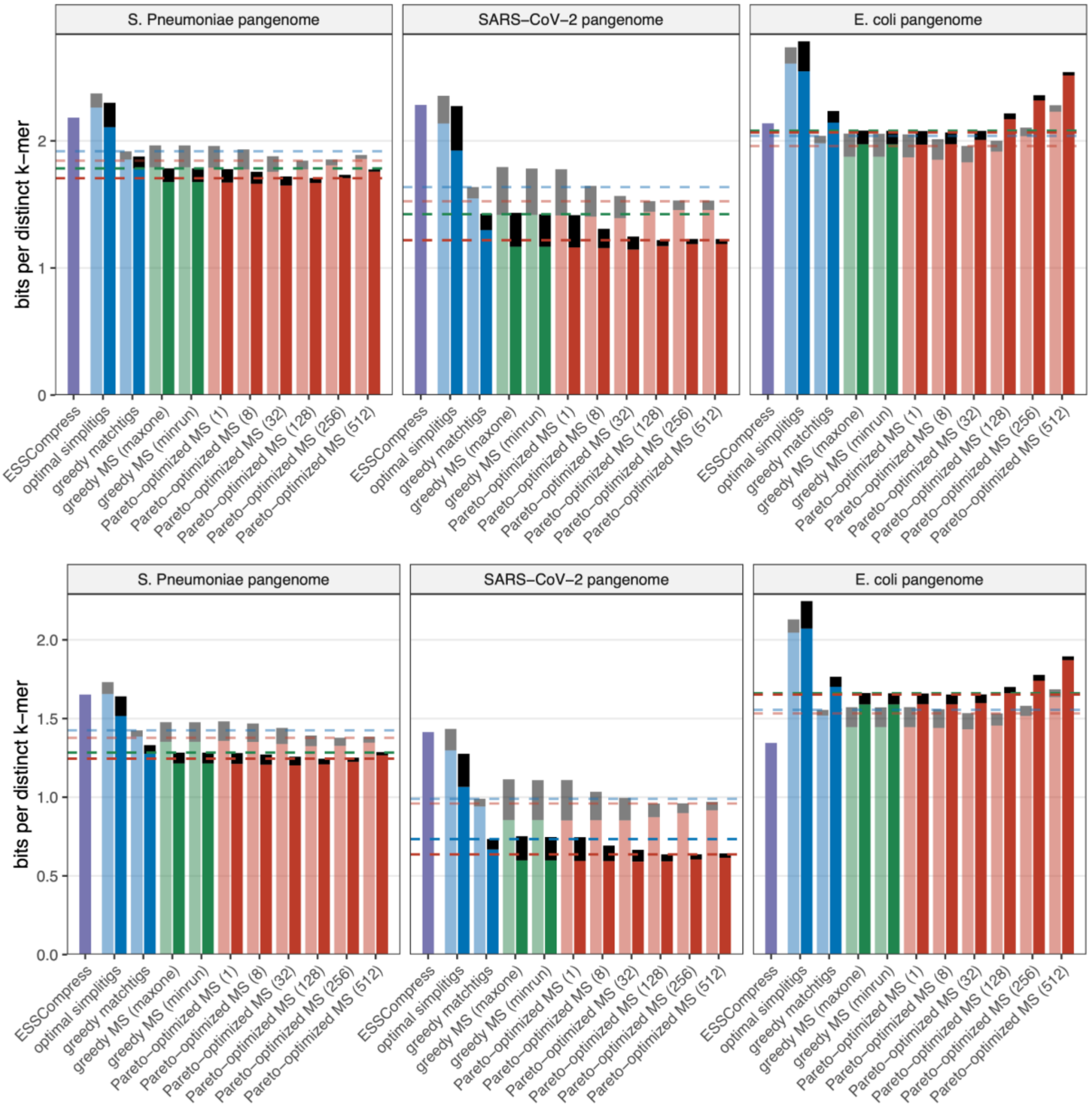

**Fig. S2. Disk compression of MS representations for selected datasets using standard compression setup for $k = 31$ (top) and $k = 63$ (bottom).**

For selected methods and parameters, we display the compressed sizes of superstrings (coloured bars starting at zero) and masks (black bars starting at the top of superstring ones). The bar for ESSCompress represents the total size of the $k$-mer set compressed with ESSCompress. Opaque bars represent compression with GeCo3, while semi-transparent bars represent compression with xz. The parameters for these tools are specified in presets 1 and 3 in **Note S3**. The green or blue dashed line represents the best compressed size of the state-of-the-art representation (masked superstrings with min-run mask optimization or matchtigs), and the red dashed line represents the best compressed size achieved using Pareto optimization. For standard compression of simplitigs and matchtigs compressed with xz, the mask has been preprocessed into a sequence of lengths of runs of ones before the compression.

### Figure S3: In-memory size with mask compression

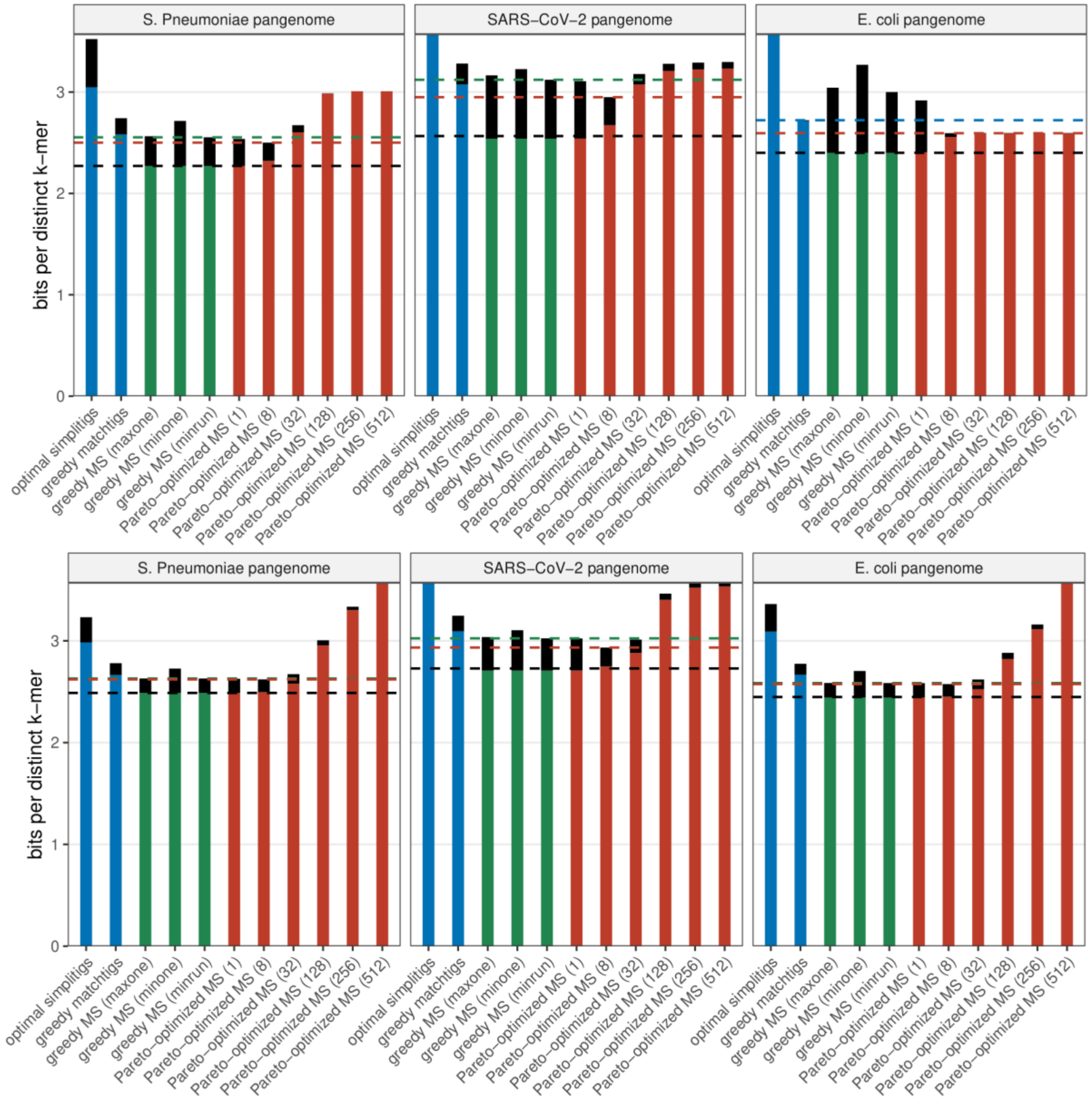

**Fig. S3. In-memory size of masked superstring representations with Elias-Fano encoded mask for selected datasets and $k = 15$ (top) and $k = 31$ (bottom).**

For selected methods and parameters, we display the in-memory sizes of computed representations, in bits per $k$-mer. Coloured bars represent the size of superstrings encoded by two bits per character, black bars (starting atop the superstring ones) represent sizes of masks compressed by the Elias-Fano encoding. For SPSS, end positions of runs are only stored for runs of ones, along with the value of $k$. For MS-based methods, end positions of runs of both ones and zeros are stored. The green or blue dashed line represents the best compressed sizes of the state-of-the-art representations, the red dashed line represents the best compressed size achieved using Pareto optimization. The black dashed line indicates a lower bound on the in-memory size, a sum of the lower bound on the superstring length and the number of bits of the Elias-Fano encoding (computed from the lower bounds on the number of runs).

### Figure S4: Distinct $k$-mers appearing in the superstring

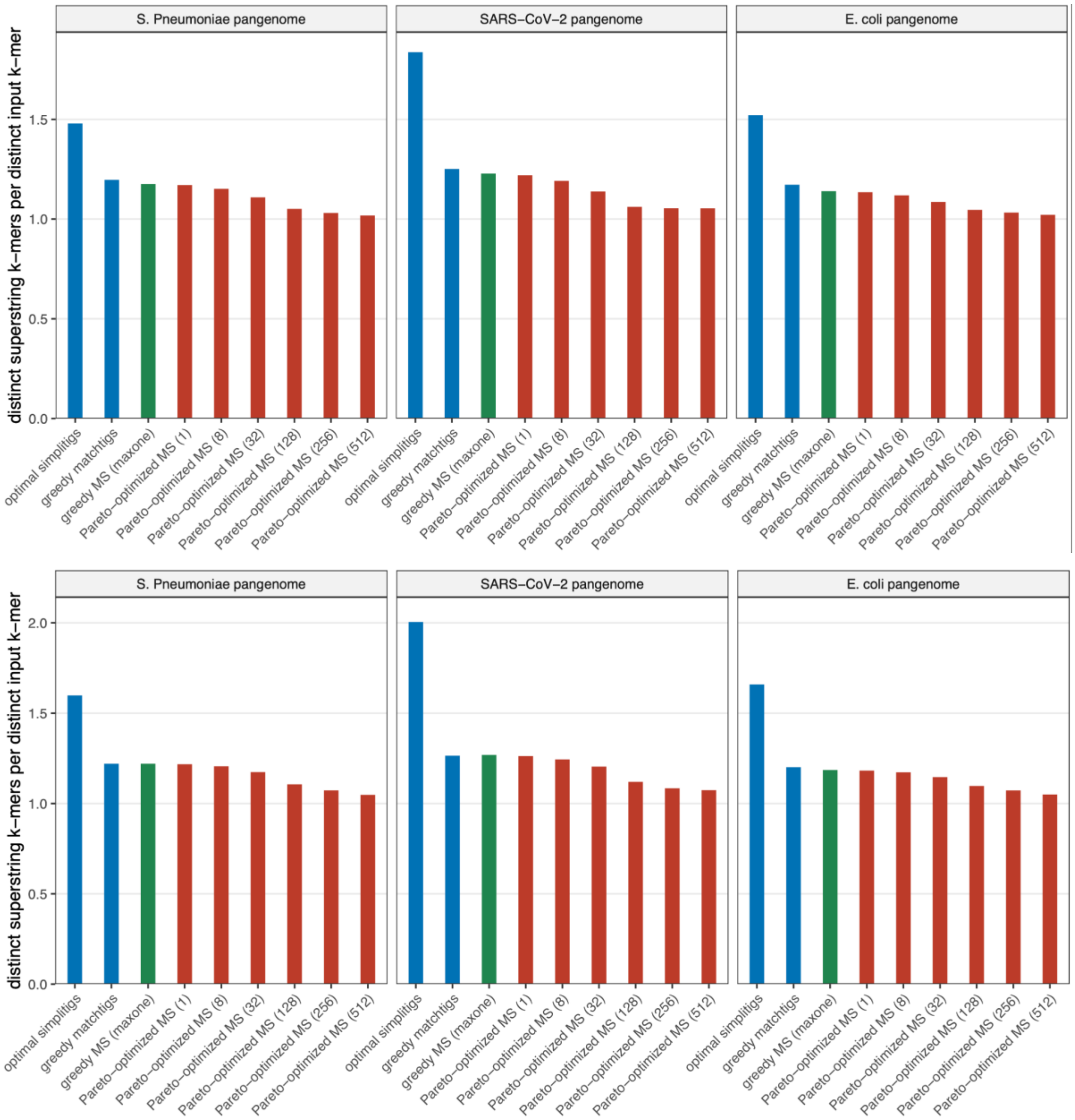

**Fig. S4. The number of distinct $k$-mers appearing in the superstring per $k$-mer from the original k-mer set, for $k = 31$ (top) and $k = 63$ (bottom).** Every original $k$-mer is masked with a 1 at least once, additional $k$-mers are always masked with zero. The trivial lower bound of 1 corresponds to only using $k$-mers from the original set in the superstring. Only distinct $k$-mers are counted, even though $k$-mers masked with both ones and zeros may appear in the superstring any number of times.

### Figure S5: Compression effectiveness of long and repetitive superstrings

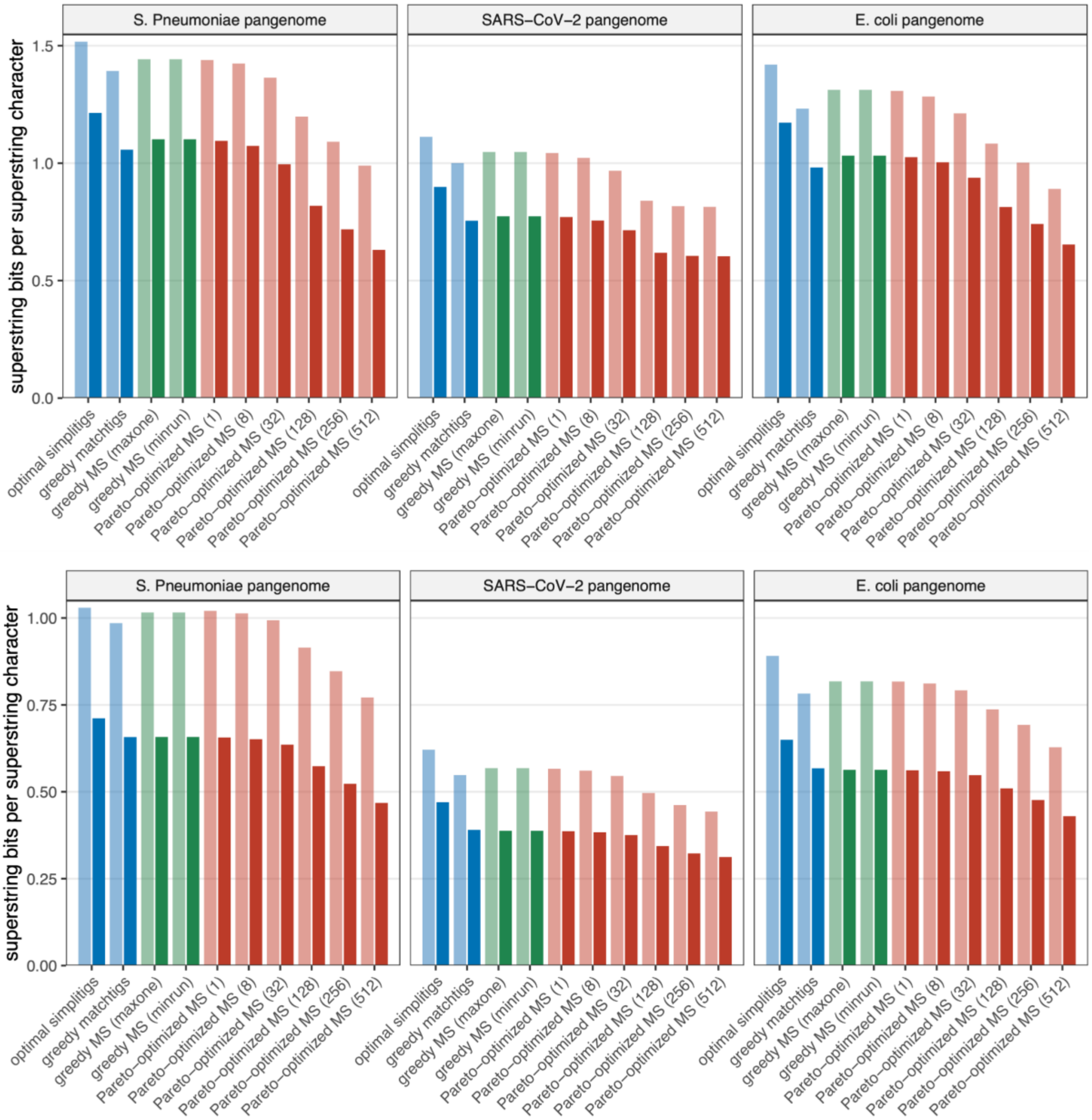

**Fig. S5. Size of compressed superstring representation per length of the superstring, in bits, for $k = 31$ (top) and $k = 63$ (bottom).** Longer superstrings reach higher repetitiveness and better compression with respect to the length. Note that the compression effectiveness increases with increasing run penalty for both neural-network-based compression (GeCo3 – opaque bars) and a classical dictionary-based compression (xz – semi-transparent bars). The length of the superstring also increases; however, the overall effect favors larger values of run penalty; see **Fig 2**.

### Supplementary tables

#### Table S1: Comparison of level penalties of MS construction methods

| **Method** | Simplitigs | Matchtigs | Greedy MS | Pareto-optimized MS |
|---|---|---|---|---|
| **Level penalties** | | | | |
| $\pi_k$ | $1$ | $1$ | $1$ | $1$ |
| $\pi_{k-1}$ | $k - 3/2$ | $2k - 3$ | $1$ | $P + 1$ |
| $\pi_{k-2}, \ldots, \pi_0$ | $-1$ | $-1$ | $1$ | $1$ |
| **Implied by the level penalties** | | | | |
| **All runs of zeros of length $k$-1** | YES | YES | NO | NO |
| **Penalty per every character** | - | $1$ | $1$ | $1$ |
| **Penalty per 1 in M** | $1$ | - | - | - |
| **Penalty per run of ones** | $1/2$ | - | - | $P$ |
| **Total penalty** | $\lvert M\rvert_1 + runs_1(M)/2$ | $\lvert S\rvert$ | $\lvert S\rvert$ | $\lvert S\rvert + P \cdot runs_1(M)$ |

An overview of level penalties which can be used to obtain a different $k$-mer representation for optimal closed covering walks in the AC automaton. Setting the penalty for upper levels to negative for simplitigs and matchtigs ensures that each run of zeros is of length $k - 1$. The penalties for additional character, for a new run and for adding $1$ into the mask are only conceptual and are implied by the setting of level penalties $\pi$. Note that the total penalty for simplitigs ensures both that they do not contain a $k$-mer more than once and that they are of minimum length.

#### Table S2: Indicative computational resources used for SARS-CoV-2 pan-genome

| | | ***k*** **= 15** | | ***k*** **= 31** | | ***k*** **= 63** | |
|---|---|---|---|---|---|---|---|
| **Method** | **Parameter** | **Time [s]** | **Mem [MB]** | **Time [s]** | **Mem {MB]** | **Time [s]** | **Mem [MB]** |
| Shortest simplitigs | | 12.5 | 1025 | 11.2 | 917 | 14.7 | 1156 |
| Greedy matchtigs | | 18.7 | 927 | 16.3 | 908 | 28.7 | 1034 |
| Greedy MS | min-one | 7.0 | 90 | 19.1 | 159 | 81.2 | 1089 |
| | max-one | 7.0 + 0.6 | 90 | 19.1 + 1.9 | 159 | 81.2 + 6.0 | 1089 |
| | min-run | 7.0 + 1.8 | 544 | 19.1 + 5.2 | 1178 | 81.2 + 17.8 | 4485 |
| Pareto--optimized MS | *1* | 22.9 | 432 | 99.8 | 1055 | 1014.1 | 2891 |
| | *7* | 27.5 | 438 | 99.6 | 1061 | 981.2 | 2899 |
| | *30* | 29.6 | 448 | 103.9 | 1073 | 905.1 | 2918 |
| | *70* | 45.9 | 451 | 113.3 | 1095 | 835.3 | 2945 |
| | *250* | 187.5 | 454 | 254.4 | 1116 | 1051.2 | 3016 |
| | *400* | 311.2 | 454 | 474.8 | 1117 | 1343.7 | 3035 |

The experiments where performance measurements were considered were run using a laptop with a 13th Gen Intel Core i7-13620H CPU with maximum frequency of 4.9 GHz and three levels of cache. For other datasets, the experiments were run using MetaCentrum grid computing infrastructure.

The SARS-CoV-2 pan-genome dataset contains 16 million genomes and 12 million 31-mers. Larger datasets require substantially more time.

### Supplementary algorithms

#### Alg S1: Pseudocode for Pareto optimization of MS

```
// Computes the Pareto-optimized MS given k-mer set K and run penalty P
function ParetoOptimization(k, K, P):
    AddReverseComplements(K)
    Sort(K)
    paths ← K
    for penalty limit from 1 to P + k:
        foreach path in paths:
            TryMergePath(path, penalty limit)
    return paths[0]

// Searches any other path that can be merged with a given path adding at most penalty limit
function TryMergePath(path, penalty limit):
    search begin ← { index: LastKmer(path), depth: k, penalty: 0 }
    RunDFS(search begin) with currently visited DFS node as index, depth, penalty:
        if depth = k and Path(index) ≠ path: // Checked using Union-find
            Merge(path, Path(index))
            return

        failure index, failure depth, penalty increase ← Rise(index, depth)
        if penalty + penalty increase ≤ penalty limit:
            ExtendDFS(failure index, failure depth, penalty + penalty increase)

        if depth < k:
            foreach leaf index in Fall(index, depth):
                ExtendDFS(leaf index, k, penalty)

// Return all leaf nodes of the subtree rooted in the current node
function Fall(index, depth):
    foreach leaf index in K from index while Prefix(index, depth) = Prefix(leaf, depth):
        yield leaf index

// Follow the failure edge from the current node, return the node found and penalty increase
function Rise(index, depth):
    for new depth in depth − 1 to 0:
        new index ← BinSearchPrefix(Suffix(index, new depth))
        if new index ≠ |K|:
            penalty increase ← depth − new depth
            if depth ≥ k − 1 and new depth < k − 1:
                penalty increase += P
            return new index, new depth, penalty increase
    return 0, ∞
```

### Supplementary notes

#### Note S1: NP-hardness of Pareto optimization

**Theorem 1 (NP-hardness of Pareto optimization).** *For any constant $P > 0$ and any value of $k > 4\, log_4(|K|) + 5$, where $K$ is a set of $k$-mers, computing an optimal MS of $K$ according to the objective function $|S| + P \cdot runs_1(M)$ is NP-hard.*

**Proof**: First, we give the proof for even values of $k$. We reduce from the NP-hard shortest superstring problem for $l$-mer sets in the bidirectional model [1], where $l = k/2$. We first transform the input $l$-mer set $L$ into the $k$-mer set $K$, solve the Pareto optimization problem for $K$ and obtain MS, which we transform into $MS_L$ of $L$ with optimal $S_L$.

We construct $K$ in the following way: For each $l$-mer $L$, we replace every character with a digram according to the mapping $\mu$: A : AC C : AT G : GC T : GT

We obtain a set of $k$-mers where all pairs of $k_i$, $k_j$ have an overlap of even length, because characters A and G only appear at odd positions and characters C and T at even positions in $k$-mers. This also holds for complements of $k$-mers and therefore works in the bidirectional model as well. Therefore, no two $k$-mers have an overlap of length $k - 1$. This means that the minimum number of runs of ones in $M$ we can obtain is $|K|$.

However, we can obtain exactly $|K|$ runs by optimally arranging the $k$-mers of $K$ (choosing the optimal permutation). If we used any $k$-mer not in $K$ or used some $k$-mer of $K$ more than once, the number of runs of ones in $M$ may only increase (if the $k$-mer is in $K$) and the length of $S$ would not decrease. Therefore, the optimal MS has exactly $|K|$ runs of ones and the optimality of MS depends only on the optimality of the length of $S$.

Solving the Pareto optimization therefore results in an optimal $S$, which can be directly transformed into the optimal $S_L$ of $L$ using the inverse mapping for each pair of characters in $S$. Therefore, the shortest superstring problem for $l$-mers (which is NP-hard for $l = k/2 > log_4 |L|$ can be solved by solving the problem of Pareto optimization of the objective function of the form $|S| + P \cdot runs_1(M)$ for $k$-mers, which means that Pareto optimization of such an objective function is also NP-hard for even values of $k$.

Next, we prove this also for odd values of $k$ by modifying the previous part of the proof. First, we choose $l$ to be $(k - 5)/4$ if $k \equiv 1 \pmod 4$ or $(k - 3)/4$ if $k \equiv 3 \pmod 4$; the other values of $k$ are already covered. Using the mapping $\mu$ from previous proof, we encode each $l$-mer $L_i$ from $L$ into $2l$-mer $L_i'$ . We then construct $k$-mers of $K$ by concatenating $L_i'$, CCC, $L_i'$ or $L_i'$, CCCCC, $L_i'$. This results in $k$-mers of length $4l + 3$ or $4l + 5$, respectively. We observe that no two $k$-mers have an overlap of length $> k/2$, which also holds in the bidirectional model. Therefore, no two $k$-mers have an overlap of length $k - 1$ and the least possible number of runs in the mask obtained by Pareto optimization is $|K|$. The rest of the proof is the same as in the previous part. The Pareto optimization problem is therefore NP-hard for all values of $l = (k - 5)/4 > log_4 |L|$, and for all $k > 4\, log_4 |K| + 5$. **Q.E.D.**

Note that the decision version of the problem is itself in NP, as the masked superstring serves as a certificate that can be validated in polynomial time. Hence, the problem is NP-complete.

### Note S2: Details of the lower bound on the number of runs

Here, we provide a proof of Lemma 1 and discuss details of computing the lower bound on the number of runs in the bidirectional model.

**Lemma 1 (unidirectional).** *Given a $k$-mer set $K$, the minimum number of runs in the mask of any MS representing $K$ in the unidirectional model is exactly equal to the minimum number of matchtigs needed to represent $K$ in the unidirectional model.*

> **Proof:** Recall that every MS corresponds to a walk in the overlap graph of $k$-mers. A new run in the mask is created whenever the walk takes an edge of overlap shorter than $k-1$, i.e., whenever it leaves its de Bruijn spanning subgraph. The walk can be split on these edges and the number of runs then exactly matches the number of these paths in the de Bruijn graph. Conversely, each path cover of de Bruijn graph can be turned into an MS with the number of runs equal to the number of paths. Hence, the minimum number of runs possible directly corresponds to the minimum number of paths covering the de Bruijn graph of $K$, i.e., to the minimum number of matchtigs representing $K$. **Q.E.D.**

Note that Lemma 1 holds also for the bidirectional model via the same reasoning. However, we do not have an efficient algorithm for computing the minimum number of matchtigs in the bidirectional model. Therefore, we relate the number of runs in the bidirectional model to the number of matchtigs in the unidirectional model for an adjusted $k$-mer set.

**Lemma S1 – bidirectional.** *Given a $k$-mer set $K$, the minimum number of runs in the mask of any MS representing $K$ in the bidirectional model is lower bounded by half of the minimum number of matchtigs needed to represent $K$ with added reverse complements in the unidirectional model.*

> **Proof:** Consider matchtigs of the smallest count possible in the bidirectional model for $K$. Let $K'$ be $K$ with added reverse complements. Observe that these matchtigs together with their reverse complements cover $K'$. Thus, they provide an upper bound on the smallest number of matchtigs for $K'$ in the unidirectional model. Therefore, the number of matchtigs of $K'$ for the unidirectional model divided by two lower-bounds the smallest number of matchtigs of $K$ in the bidirectional model. **Q.E.D.**

Note that the resulting lower bound for the bidirectional model is not generally tight. In the bidirectional model, for every walk $w$ in the de Bruijn graph of $K$, there exists a reverse complementary walk $w'$ that only differs in the direction. It may happen that $w$ and $w'$ overlap in every $k$-mer, and we call such $w$ a *self-complementary* walk. The resulting lower bound for the bidirectional model is not tight due to the existence of self-complementary walks, which may be covered by a single matchtig in both models, but are counted only as a half in the result; however, the result is still a lower bound. Also, note that the difference between the lower bound computed for the bidirectional model using this approach and the optimal number of runs in the mask of any MS in the bidirectional model is upper bounded by the number of strongly connected components in the de Bruijn graph of $K$ that contain both the $k$-mer and its reverse complement. In our experiments, this number ranged from *1* for smaller datasets to *45* for larger ones. In the implementation, we use a more complex approach, handling several edge cases to further tighten the bound in practice.

### Note S3: Details of experimental evaluation

Simplitigs and greedy matchtigs were computed using GGCAT [2] (v2.0.0).

Greedy masked superstrings were computed and masks optimized using Kmercamel [1] (v2.2.0), which was also used for computing the lower bounds of the superstring length.

Lower bounds for the number of runs in the mask and masked superstrings produced by iterative deepening search for Pareto optimization were computed using our own implementation, using basic functions for $k$-mer manipulation from Kmercamel.

Data were compressed using xz v5.8.1, bzip2 v1.0.8, and GeCo3 [3] v1.0, with four presets:

1. Standard compression using standard tools: superstring and mask: xz -9. For simplitigs and matchtigs, the mask was first encoded as a sequence of lengths of runs of ones.
2. Standard compression using GeCo3: superstring and mask: GeCo3 -l5.
3. Maximal compression using standard tools: superstring: xz -e --lzma2=preset=9,dict=1500MiB,nice=250, mask: bzip2 --best. For simplitigs and matchtigs, the mask was first encoded as a sequence of lengths of runs of ones.
4. Maximal compression using GeCo3: superstring and mask: GeCo3 -lr 0.005 -hs 160 -tm 1:1:1:0:0.6/0:0:0 -tm 1:1:0:0:0.6/0:0:0 -tm 2:1:2:0:0.90/0:0:0 -tm 2:1:1:0:0.8/0:0:0 -tm 3:1:0:0:0.8/0:0:0 -tm 4:1:0:0:0.8/0:0:0 -tm 5:1:0:0:0.8/0:0:0 -tm 6:1:0:0:0.8/0:0:0 -tm 7:1:1:0:0.7/0:0:0 -tm 8:1:0:0:0.85/0:0:0 -tm 9:1:1:0:0.88/0:0:0 -tm 11:10:2:0:0.9/0:0:0 -tm 11:10:0:0:0.88/0:0:0 -tm 12:20:1:0:0.88/0:0:0 -tm 14:50:1:1:0.89/1:10:0.89 -tm 17:2000:1:10:0.88/2:50:0.88 -tm 20:1200:1:160:0.88/3:15:0.88 (the best-performing preset from the human genome compression experiment available at https://github.com/cobilab/HumanGenome; Run39.sh; retrieved on March 11, 2026).

Note that even though the GeCo3 tool uses a neural network to compress the sequences, the neural network is not pretrained on our datasets and uses only the preset settings and the sequences being compressed as inputs.

For the compression baseline, we used ESSCompress [4] (v3.1) with default settings.

To check that our methods work correctly in combination with the neural-network-based compression, we also included the decompression step (using GeDe3 [3]) in the pipeline. The number of distinct $k$-mers was computed (using jellyfish) for the initial dataset, the decompressed representation (converted to SPSS) and the union of both, to ensure the decompressed representation contains exactly the same $k$-mers as the dataset before compression, i.e., that the compression is lossless.

The source code and pipelines for experiments are provided in the **Supplementary repository** (https://github.com/Jajopi/ms-pareto-optimization-supplement).
